# Autophagy inhibition rescues structural and functional defects caused by the loss of mitochondrial chaperone *Hsc70-5/mortalin* in *Drosophila*

**DOI:** 10.1101/2020.09.17.302141

**Authors:** Jun-yi Zhu, Shabab B. Hannan, Nina M. Dräger, Natalia Vereshchagina, Ann-Christin Krahl, Yulong Fu, Christopher J.H. Elliott, Zhe Han, Thomas R. Jahn, Tobias M. Rasse

**Affiliations:** Hertie Institute for Clinical Brain Research, University of Tübingen, Otfried-Müller-Str. 27, 72076 Tübingen, Germany; Center for Genetic Medicine Research, Children’s National Medical Center, Washington, DC 20010, USA; Schaller Research Group at the University of Heidelberg and DKFZ, Proteostasis in Neurodegenerative Disease (B180), German Cancer Research Center, Im Neuenheimer Feld, 69120 Heidelberg, Germany; Department of Biology, University of York, York, YO1 5DD, UK; Max Planck Institute for Heart and Lung Research, Ludwigstr. 43, 61231 Bad Nauheim, Germany; AbbVie Deutschland GmbH, Knollstr., 67061 Ludwigshafen, Germany

**Keywords:** *Atg1*, *Hsc70-5*, microtubule, mitochondria, mitophagy, *mortalin*, rapamycin, synapse

## Abstract

We investigate in larval and adult *Drosophila* models whether loss of the mitochondrial chaperone *Hsc70-5*/*mortalin* is sufficient to cause pathological alterations commonly observed in Parkinson disease. At affected larval neuromuscular junctions, no effects on terminal size, bouton size or number, synapse size, or number were observed, suggesting that we study an early stage of pathogenesis. At this stage, we noted a loss of synaptic vesicle proteins and active zone components, delayed synapse maturation, reduced evoked and spontaneous excitatory junctional potentials, increased synaptic fatigue, and cytoskeleton rearrangements. The adult model displays ATP depletion, altered body posture, and susceptibility to heat-induced paralysis. Adult phenotypes could be suppressed by knockdown of *DJ-1b, LRRK, p50, p150, Atg1, Atg101, Atg5, Atg7*, and *Atg12*. The knockdown of components of the autophagy machinery or overexpression of human *mortalin* broadly rescued larval and adult phenotypes, while disease-associated *HSPA9* variants did not. Overexpression of *Pink1* or promotion of autophagy exacerbated defects.

## Introduction

Parkinson disease (PD), the most prevalent movement disorder and the second most prevalent neurodegenerative disease, is characterized by resting tremor, stiffness, and slowness of movement. Analysis of genetic and environmental factors contributing to PD suggests that impairments in mitochondrial function, lysosomal degradation pathways, and synaptic transmission are of central importance to pathogenesis and progression of PD [1-4]. Due to their complex morphology and high-energy demands, neurons in the adult brain are particularly susceptible to dysregulation of mitochondrial quality control systems. These quality control systems operate at different levels protecting cells and tissues from dysfunctional mitochondria. While a significant proportion of PD cases are sporadic, mutations in at least 11 genes have been implicated in monogenic typical or atypical forms of parkinsonism [3], providing crucial insights into the cellular and molecular pathways involved in PD. In all cases, the detailed molecular mechanisms leading to disease development have not been fully elucidated. However, a number of PD-associated genes have been implicated in mitochondrial function, altered mitochondrial dynamics, or the accumulation of dysfunctional mitochondria, which are characteristic features of PD [6-11]. For example, the PD-associated genes *PTEN-induced kinase 1* (*Pink1*) and *parkin* are important regulators of mitochondrial quality control and mitophagy, a form of macroautophagy [6-8]. Mitochondrial dysfunctions have also been implicated in various other neurodegenerative diseases such as Alzheimer disease where mitochondria are key targets of both Aβ42 and tau toxicity [12].

We have previously addressed the importance of mitochondrial quality control and selective vulnerability of dopaminergic neurons by generating a model for loss-of-function of *Drosophila Hsc70-5/mortalin* (referred to as *Hsc70-5* or *mortalin* hereafter) [13]. Enhanced mitophagy was identified as an early pathological feature caused by knockdown in a presymptomatic model of loss of *Hsc70-5* function [13] prior to the emergence of locomotion defects. Consistent with the upregulation of mitophagy upon loss of *Hsc70-5* function in a *Drosophila*, human fibroblasts obtained from a carrier of the PD-associated A476T Mortalin variant exhibit increased colocalization of mitochondria with autophagosomes.

In the current study, we investigate in detail the consequences of neuronal loss of *Hsc70-5/mortalin* function *in vivo* and examine epistatic interactions in this genetic background.

## Results

### Impairment of larval locomotion upon loss of *Hsc70-5/mortalin* function

We have previously shown that strong pan-neuronal expression (*elav-Gal4, 29°C*) of the RNAi-construct *mort*^*KK*^ results in lethality at the late L3 larval stage [14]. In the current study, we sought to investigate cellular and functional perturbations that occur early in pathogenesis. We thus reduced silencing potency by raising larvae at 25°C, which delayed lethality to the pupal stage (elav>*mort*^KK^ and elav>*mort*^GD^). elav>*mort*^*KK*^ larvae were sluggish compared to size-matched elav>*mort*^GD^ and control larvae. Their crawling velocity was reduced and the righting reflex delayed (Figure 1A). elav>*mort*^*GD*^ larvae were indistinguishable from controls in their righting ability. Hence we referred to them as presymptomatic and elav>*mort*^KK^ larvae as symptomatic (Table S1). *Hsc70-5* is an important regulator of mitochondrial function [13,15-17]. Thus, we investigated putative mitochondrial impairments in presymptomatic and symptomatic larvae. Pan-neuronal *Hsc70-5* knockdown resulted in severe reductions in mitochondrial mass at neuromuscular junctions (NMJs) of symptomatic (elav>*mort*^*KK*^, 25°C) but not presymptomatic (elav>*mort*^*GD*^, 25°C) larvae compared to control (Figure 1B). Symptomatic larvae showed a reduction in mitochondrial number and size, and consequently, a decrease in mitochondria area fraction at the NMJ (Figure S1). Muscle length, NMJ area, and bouton numbers were unaltered (Figure S2) in both presymptomatic and symptomatic larvae. This suggests that elav>*mort*^*KK*^ larvae represent an early stage in disease progression characterized by the absence of gross neurodevelopmental or neurodegeneration phenotypes.

**Figure 1.**
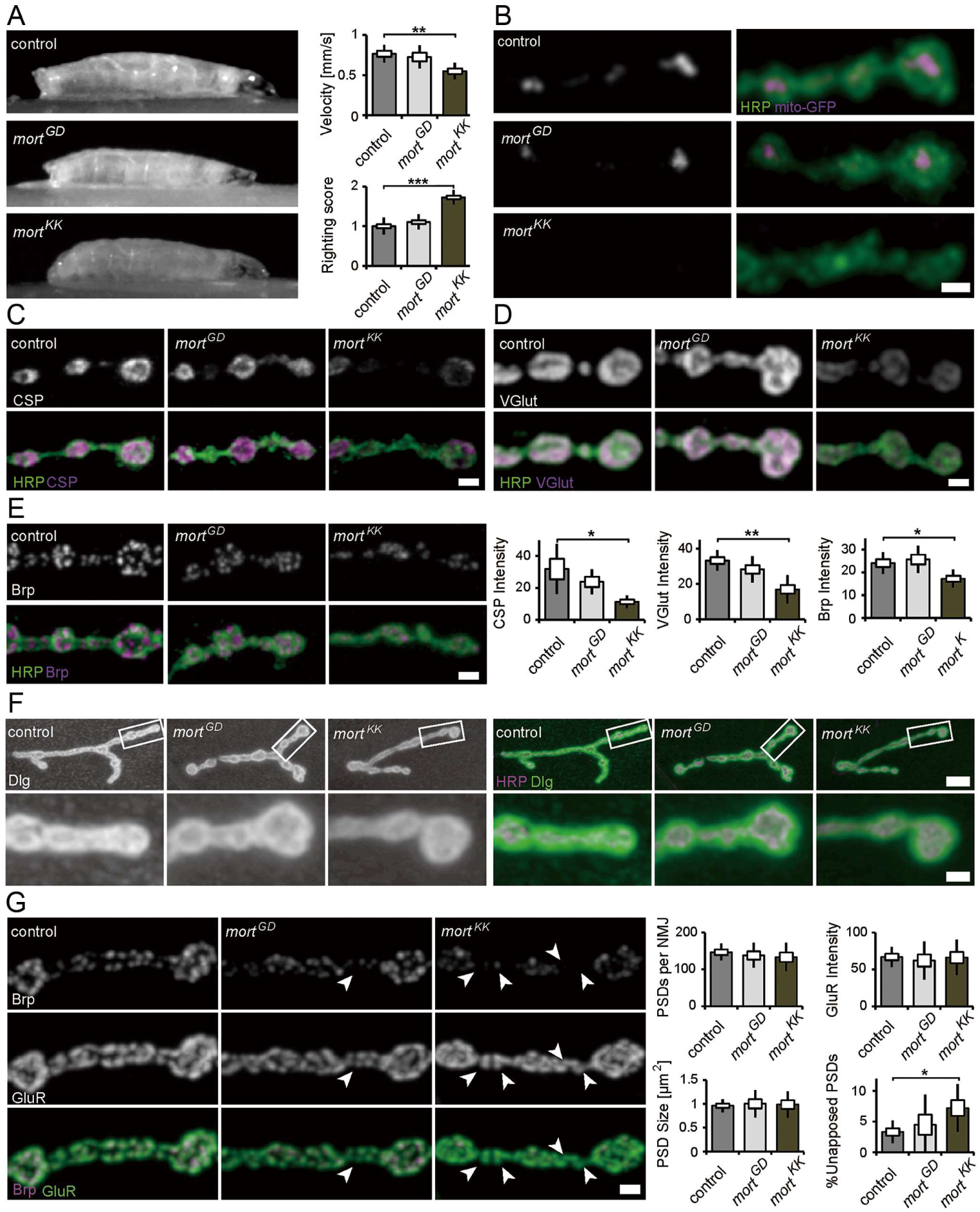
Characterization of the neuromuscular junction in presymptomatic and symptomatic larval. (**A**) Average larval crawling velocity and righting reflex at L3 stage. (**B**) Confocal images of larval NMJ expressing mito-GFP (magenta) and labeled with HRP (green). Scale bar: 2 μm. (**C-E**) Representative synaptic boutons from NMJs of control, elav>*mort*^*GD*^ and elav>*mort*^*KK*^ larvae. Anti-HRP staining (C-E, green) was used to visualize synaptic membranes. Synaptic vesicle proteins CSP (**C**) and VGlut (**D**) and active zone component Bruchpilot (Brp) (**E**) are shown in magenta. Scale bar: 2 μm. (**F**) Confocal images of larval NMJ labeled with HRP (magenta) and *Drosophila* discs large (Dlg; green) used to visualize the sub-synaptic reticulum (SSR). Scale bar: 5 μm, enlargement: 2 μm. (**G**) Confocal images of larval NMJ labeled with Brp (magenta) and GluR (green) were used to visualize postsynaptic densities (PSDs). Arrowheads pointing out presynaptic Brp labels were not detected in PSDs. Scale bar: 2 μm. The standard error of mean (SEM) and standard deviation (SD) are shown as a box and a black line, respectively. * p<0.05, ** p<0.01, *** p<0.001.

### Detection of early synaptic changes due to loss of *Hsc70-5/mortalin* function

To further investigate structural changes in response to loss of *Hsc70-5* function, we stained for synaptic vesicles (SVs) and active zone (Az) markers. To this aim we performed confocal imaging on fixed larval NMJs. Analysis and quantification revealed that SV proteins cysteine-string protein (CSP, Figure 1C) and the vesicular glutamate transporter (VGlut, Figure 1D) were reduced at NMJs of symptomatic larvae compared to control larvae. A similar reduction was observed for the central organizing component of Azs, Bruchpilot (Brp, Figure 1E). We observed no significant differences in presymptomatic larvae for SV proteins or Az components compared to control (Figure 1C-E).

We proceeded to investigate synapse maturation by analyzing subsynaptic reticulum (SSR) and postsynaptic density (PSD) maturation. The SSR, a complex system of membrane tubules and lamellae, is formed by the invagination of postsynaptic membranes following presynaptic innervation. Thus, an increased percentage of terminal synaptic boutons devoid of SSR can be used as a marker for delayed synapse terminal maturation [18,19]. To examine SSR folding and morphogenesis, we labeled NMJs with antibodies against Dlg and the neuronal marker HRP. Dlg reliably surrounds terminal boutons in all genotypes examined, suggesting that *Hsc70-5* knockdown does not affect SSR morphogenesis (Figure 1F). Furthermore, quantification of the number, intensity, and size of glutamate receptor cluster per synapse (Figure 1G) revealed no postsynaptic phenotypes.

We next assessed defects in presynaptic maturation by quantifying the percentage of PSDs unopposed by Azs. To this aim, we analyzed the fraction of PSDs apposed by Az component Brp that reliably localizes to mature synapses. Az apposed to newly forming PSDs usually recruit detectable Brp puncta within 2-4 h of their formation and, more than 95% of all glutamatergic synapses are apposed by Brp [20]. A higher percentage of Brp-negative synapses is indicative of neurodegenerative or neurodevelopmental defects [21]. elav>*mort*^*KK*^ larvae displayed defects in the apposition of PSDs and Azs. Twice as many unaposed, putatively immature synapses were detected. This defect might be caused by impairments in presynaptic maturation or stabilization (Figure 1G).

Defects in MT cytoskeletal organization have been associated with PD models [22,23] in particular, and with loss of synaptic mitochondria in general [24,25]. The *Drosophila* homolog of the *leucine-rich repeat kinase 2(LRRK2)*, linked to familial and sporadic PD, controls MT stability at *Drosophila* NMJs by suppressing Futsch function [23]. Additionally, a decreased number of MT loops, which is indicative of stable MT, has been observed [24]. In this study fewer MT loops were detected in terminal boutons at the NMJs of elav>*mort*^*KK*^ larvae (Figure 2A, Figure S3). In addition to defective MT morphology, we observed a reduction in the percentage of boutons connected to the stable MT network (Figure S3). This might impair long-range intracellular transport. Next, we employed the synaptic footprint assay and examined neuronal membranes by labeling HRP. Synaptic footprints are biomarkers for late-stage synapse retraction [26], while HRP inhomogeneity has been associated with early-stage synapse disassembly [25]. We visualized the presynaptic compartment by labeling for the membrane marker HRP and the SV marker VGlut. Neither synaptic footprints nor HRP inhomogeneity was detected in response to *Hsc70-5* knockdown (Figure 2B) suggesting that elav>*mort*^*KK*^ larvae represent an early stage of pathology with no evidence of synapse dismantlement.

**Figure 2.**
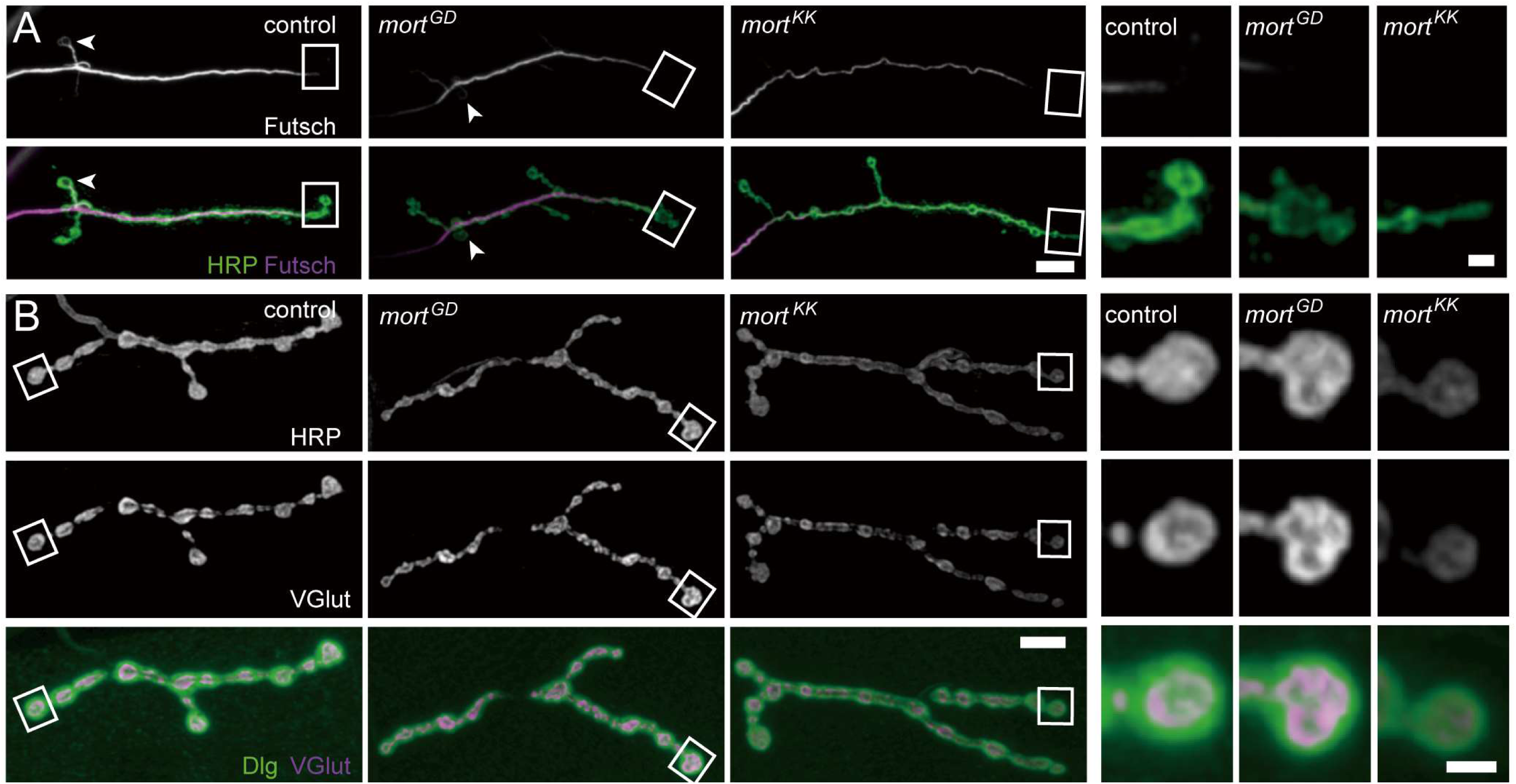
Loss of *Hsc70-5/mortalin* function is associated with MT cytoskeleton defects. (**A**) Confocal images of larval NMJ labeled with HRP (green) and Futsch (MT cytoskeleton; magenta). Scale bar: 5 μm, enlargement: 2 μm. Arrowheads pointing at NMJ loops. (**B**) Confocal images of larval NMJ labeled with HRP (gray), VGlut (gray and magenta), and Dlg (green). Scale bar: 10 μm, enlargement: 2 μm.

### Functional consequences of loss of *Hsc70-5/mortalin* function

Next, we investigated whether the aforementioned morphological changes affect synaptic function. To examine whether symptomatic elav>*mort*^KK^ larvae have impaired synaptic transmission; we recorded evoked excitatory junctional potentials (eEJPs) by using current-clamp recording at NMJ from muscle 6 in segment A5. The amplitude of eEJPs in elav>*mort*^KK^ was reduced compared to control (Figure 3A). We also recorded spontaneous events (Figure 3B and C) and the response of nerve terminals during and following high-frequency stimulations (Figure 3D-F). We noted a decrease in the amplitude of miniature excitatory junctional potential (mEJP) and an increase in their frequency in elav>*mort*^KK^ larvae compared to control. The ratio of eEJP to mEJP or quantal content (Figure 3C) was reduced in mutants compared to control. 10Hz stimulation at mutant terminals caused no changes in EJP amplitude (Figure 3D and E) during a 10Hz stimulation paradigm. However, a drastic time-dependent increase in failure in mutant compared to control (Figure 3D and F) was observed.

**Figure 3.**
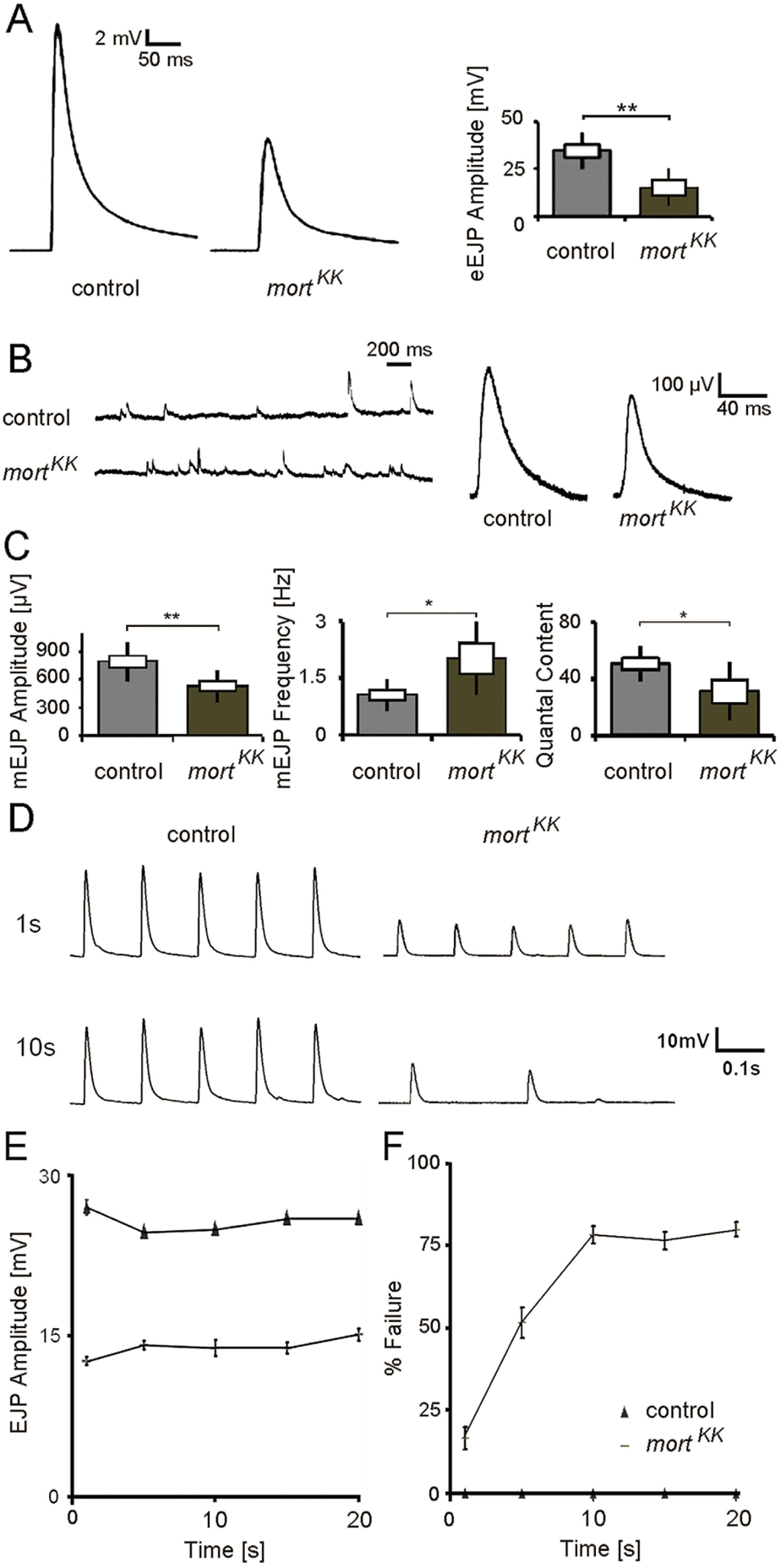
Electrophysiological characterization of symptomatic larvae. (**A**) Representative traces of eEJPs from control and *elav>mort*^*KK*^ larvae and quantification of eEJP amplitudes. Stimulation artifacts were removed from eEJP traces. (**B**) Representative mEJP recording in control and *elav>mort*^*KK*^ larvae. Quantification of (**C**) mEJP amplitude, mEJP frequency and quantal content in control and *elav>mort*^*KK*^ larvae. (**D**) Representative traces from 10Hz stimulation displaying five successive synaptic responses from each genotype after 1 and 10s stimulation. Stimulation artifacts were removed for demonstration. (**E**) Quantification of eEJP amplitude in response to 10 Hz stimulation in control and *elav>mort*^*KK*^ larvae. (**F**) Quantification of % failure of eEJP in response to 10 Hz stimulation in control and *elav>mort*^*KK*^ larvae. * p<0.05, ** p<0.01.

### Overexpression of *Drosophila* and human *mortalin* rescued *Hsc70-5/mortalin*

RNAi-induced defects

Mortalin consists of two principal domains: an ATPase binding domain and a substrate-binding domain. Rare variants R126W, A476T and P509S were identified in PD patients [17,27,28]. R126W localizes to the ATPase domain and, A476T and P509S respectively, to the substrate-binding domain of Mortalin [17,27,28]. To investigate the functional relevance of these sequence variants, we performed genetic rescue experiments in the elav>*mort*^*KK*^ background using *Drosophila* UAS*-Hsc70-5*. Pan-neuronal expression (elav-Gal4) of *mort*^*KK*^ alone or in combination with a second UAS construct expressing *lacZ* at 25°C -to control for Gal4 dilution - caused lethality at the pupal stage (Figure 4A). Co-overexpression of *Hsc70-5* with *mort*^*KK*^ restored adult viability (Figure 4A). Since human and *Drosophila mortalin* (*HSPA9* and *Hsc70-5*) have high sequence similarity [14], we examined if expression of *Hsc70-5*/*HSPA9* can rescue pupal lethality. Overexpression of *HSPA9* rescued pupal lethality, suggesting functional homology between the *Drosophila* and human *mortalin* (*Hsc70-5* and *HSPA9*) (Figure 4A). All investigated sequence variants, R126W, A476T and P509S (*HSPA9*^A476T^, *HSPA9*^P509S^ and *HSPA9*^R126W^ respectively) [17,27,28] failed to rescue pupal lethality (Figure 4A). Overexpression of either *Hsc70-5* or *HSPA9* concomitantly with *mort*^*KK*^ successfully rescued righting reflex and lowered righting time to control levels (Figure 4B). In contrast, *HSPA9*^R126W^, *HSPA9*^A476T,^ and *HSPA9*^P509S^ were unable to revert the righting defect (Figure 4B). Next, we co-overexpressed *Hsc70-5* or *HSPA9* with *mort*^*KK*^ and performed mitochondrial morphological analysis at the larval NMJ. Overexpression of *Hsc70-5* or *HSPA9* rescued number, area fraction of mitochondria at the NMJ. Moreover, normal mitochondrial size and morphology were restored. In contrast, *HSPA9*^R126W^, *HSPA9*^A476T,^ and *HSPA9*^P509S^ failed to rescue these phenotypes (Figure 4C, and C’). These results suggest that *Hsc70-5* or *HSPA9* rescued pupal lethality and larval locomotion defects by restoring *mortalin* function.

**Figure 4.**
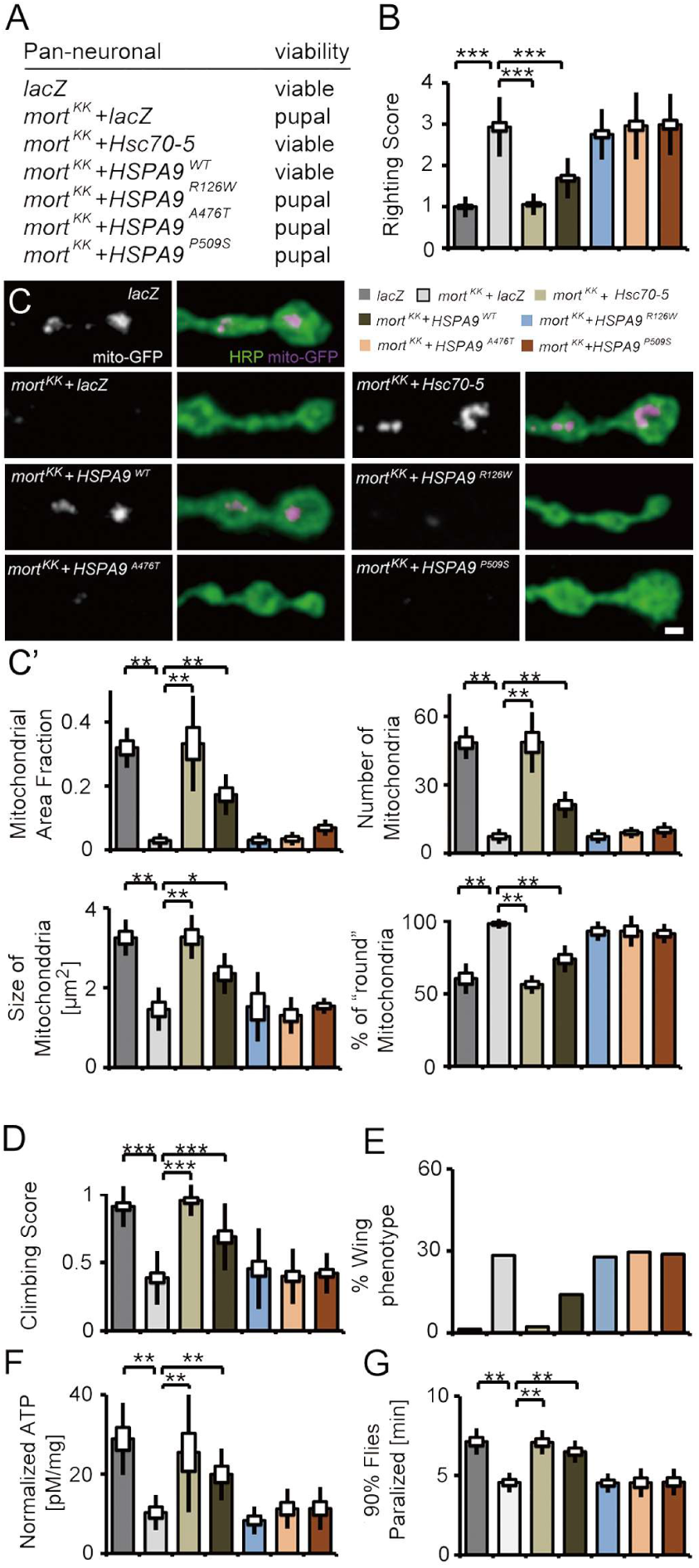
*Drosophila Hsc70-5* and human *HSPA9* but not disease variants rescued *Hsc70-5/mortalin* knockdown phenotypes. (**A**) *Hsc70-5* knockdown in combination with ectopic *lacZ* expression to control Gal4 dilution at 25°C caused pupal lethality. Overexpression of *Drosophila Hsc70-5* and human *HSPA9*^*WT*^ in the elav>*mort*^KK^ background unlike *HSPA9*^*R126W*^, *HSPA9*^*A476T*^ and *HSPA9*^*P509S*^ variants rescued pupal lethality. (**B**) Average larval righting reflex at L3 stage with overexpression of *Hsc70-5, HSPA9*^*WT*^, *HSPA9*^*R126W*^, *HSPA9*^*A476T*^ and *HSPA9*^*P509S*^ in the elav>*mort*^KK^ background. (**C**) Confocal images of larval NMJ labeled with HRP (green) and mito-GFP (magenta), and (**C’**) quantification of mitochondrial parameters. Scale bar: 2 μm. (**D**) Climbing ability of 4-day-old flies, (E) Percentage of flies with defective wing phenotype, and (**F**) ATP levels in fly heads after expressing *Hsc70-5, HSPA9*^*WT*^, *HSPA9*^*R126W*^, *HSPA9*^*A476T*^ and *HSPA9*^*P509S*^ in the *elav>mort*^*KK*^ background. (**G**) *Hsc70-5* knockdown accelerated heat-shock induced paralysis in flies at 39.5°C. The ectopic coexpression of *Hsc70-5* and *HSPA9* unlike *HSPA9*^*R126W*^, *HSPA9*^*A476T*^ and *HSPA9*^*P509S*^ rescued this defect. * p<0.05, ** p<0.01, *** p<0.001.

We utilized the Gal4/Gal80 system (elav>*mort*^*KK*^,tub-GAL80^ts^) to achieve late-onset conditional knockdown. Raising larvae at 18°C before transferring them to 25°C for Gal4 expression 5 days after egg laying (AEL) prevent pupal lethality. We thus could analyze behavioral defects in flies. 4-days post-eclosion, elav>*mort*^*KK*^,tub-GAL80^ts^ flies exhibited severe climbing defects and abnormal wing posture (Figure 4D and E). We referred to these flies as symptomatic (Table S1). Importantly, late-onset knockdown of *Hsc70-5* caused a reduction of ATP levels in fly heads (Figure 4F). Overexpression of either *Hsc70-5* or *HSPA9* in the elav>*mort*^*KK*^,tub-GAL80^ts^ background improved climbing ability, wing posture, and restored ATP levels in symptomatic model (Figure 4D-F). Overexpression of *HSPA9*^R126W^, *HSPA9*^A476T^ and *HSPA9*^P509S^ variants did not rescue any of the aforementioned defects (Figure 4D-F). ATP availability is crucial for the regulation of intracellular calcium at the presynapse, in particular upon exposure to temperature-dependent cellular stress [29-31]. Furthermore, ATP is required for reversal of membrane potential following depolarization and propagation of action potentials along axons [32]. Flies with mutations in mitochondrial protein Cytochrome c Oxidase have been reported to suffer from temperature-induced paralysis [33]. We assayed flies at 39.5°C. elav>*mort*^*KK*^,tub-GAL80^ts^ flies paralyzed faster than control flies (Figure 4G). This defect was rescued by overexpression of either *Hsc70-5* or *HSPA9* but not *HSPA9*^R126W^, *HSPA9*^A476T^ or *HSPA9*^P509S^ variants (Figure 4G).

### Downregulation of autophagy was protective

ATP and proper mitochondrial function are important maintain synaptic transmission at increased temperatures [29]. Mutants with compromised mitochondrial function paralyze faster compared to controls. We thus performed a candidate-based screen to identify modifiers of *Hsc70-5* loss-of-function (Figure 5A). Downregulation of synaptic proteins involved in neurotransmission and endocytic machinery enhanced heat-stress induced paralysis observed upon loss of *Hsc70-5*. Knockdown of Parkinson disease associated *DJ-1b* and *LRRK*, as well as components of retrograde transport machinery, were suppressors of *Hsc70-5* knockdown induced paralysis.

**Figure 5.**
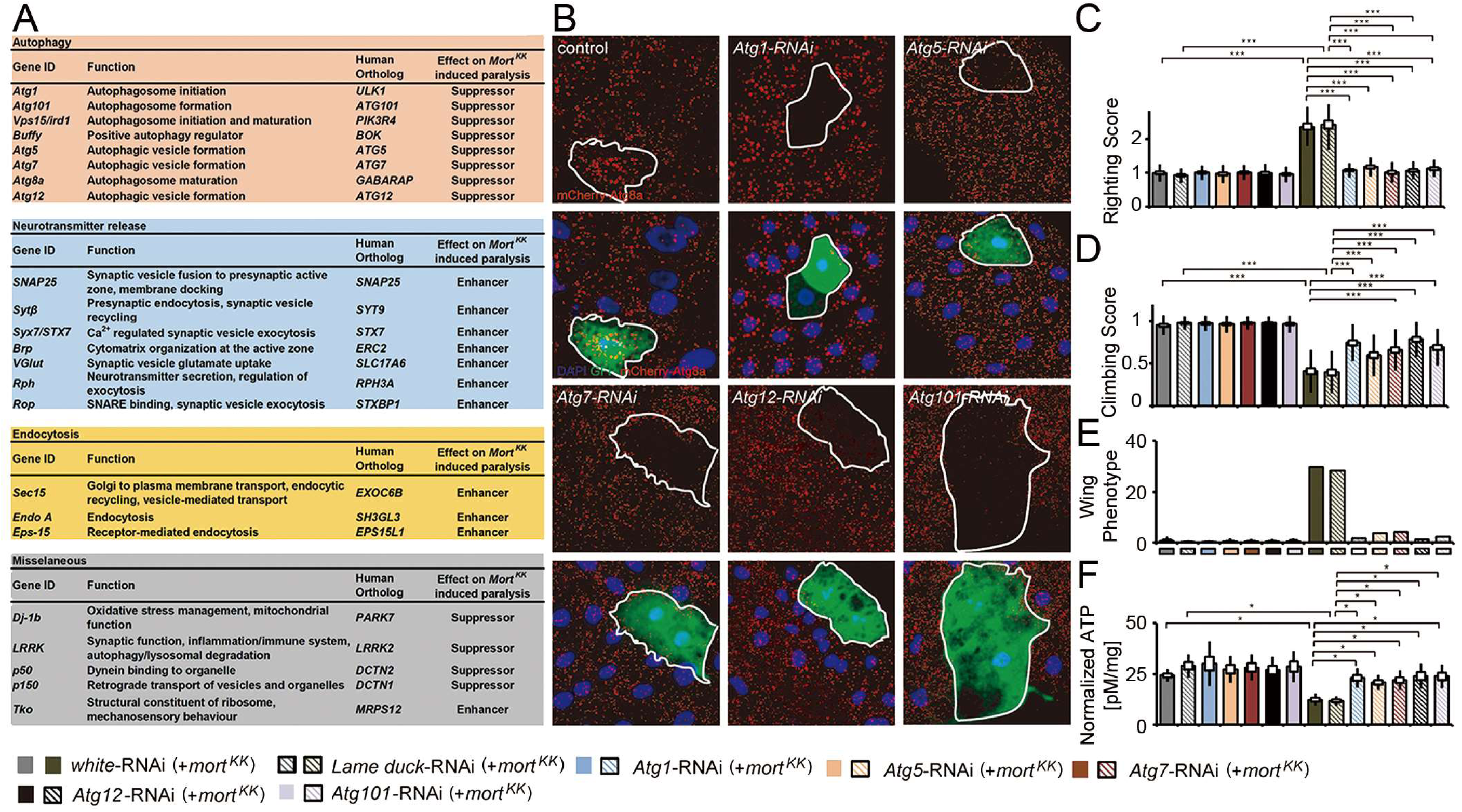
A genetic screen identified autophagy-related proteins as modifiers of *Hsc70-5/mortalin* knockdown phenotypes. (**A**) A genetic screen identified several modifiers that modified *Hsc70-5* knockdown induced phenotypes. (**B**) Increase in mCherry.*Atg8a* puncta in the larval fat body following starvation. There was no difference in the expression of mCherry.*Atg8a* between GFP +ve and -ve cells in control overexpressing *lacZ* (in GFP +ve cell). Expression of RNAi against autophagy-related genes *(Atg1, Atg101 Atg5, Atg7*, and *Atg12)* inhibited starvation-induced autophagy in GFP +ve cells compared to cells in the vicinity. Scale bar: 15 μm. (**C**) Righting reflex in larvae, (**D**) Climbing ability of 4-day-old flies, (**E**) Percentage of flies with defective wing phenotype, and (**F**) ATP levels in fly heads after pan-neuronal knockdown of *Atg1, Atg101 Atg5, Atg7*, and *Atg12* in control and the elav>*mort*^*KK*^,tub-GAL80^ts^ background. * p<0.05, *** p<0.001.

We identified five members of the autophagy machinery *Atg1, Atg5, Atg7, Atg12, and Atg101* as suppressors of *Hsc70-5* loss-of-function induced paralysis (Figure 5A). To verify the RNAi efficiency of modifiers isolated in the screen, we tested the *Atg1, Atg5, Atg7, Atg12, and Atg101* RNAi strains for their ability to inhibit starvation-induced autophagy (Figure 5B). Overexpression of RNAi against *Atg1, Atg5, Atg7, Atg12, and Atg101* in the larval fat body using a mosaic system [34,35] for genetic analysis in cells expressing GFP completely suppressed starvation-induced autophagy (Figure 5B). After validation of these constructs, we tested if knockdown of autophagy-related genes could ameliorate phenotype caused by loss of *Hsc70-5*. We used UAS-RNAi strains against *Lame duck* and *white* with *mort*^*KK*^ (*mort*^*KK*^+*white-RNAi* and *mort*^*KK*^+*Lame duck-RNAi*) as controls to compare larvae bearing *mort*^*KK*^ in combination with a second UAS site expressing RNAi against autophagic components. Pan-neuronal expression of RNAi strains targeting components of the autophagic machinery, *Atg1, Atg5, Atg7, Atg12, and Atg101* in the elav>*mort*^*KK*^ background restored the righting reflex to control levels (Figure 5C). Next, we extended the analysis to flies. As previously demonstrated 4-day-old elav>*mort*^*KK*^,tub-GAL80^ts^ symptomatic flies displayed severe defects in climbing and wing posture (Figure 5D and E) [14]. Knockdown of autophagic components reversed impairments in locomotion, wing posture and ATP levels (Figure 5 D-F) induced by *Hsc70-5* knockdown in young flies. Knockdown of autophagy genes alone did not reveal any significant differences compared to controls (Figure 5C-F). These results suggest that inhibiting autophagy is sufficient to rescue *Hsc70-5* knockdown associated impairments in larval and adult symptomatic models.

### Autophagy induction in symptomatic flies exacerbated *Hsc70-5/mortlain* knockdown induced defects

Loss of *Hsc70-5* function has been associated with increased mitophagy in *Drosophila* and human fibroblasts [14]. Consistent with a protective role for autophagy, *Parkin* overexpression is sufficient to rescue alterations in mitochondrial morphology and increased apoptosis [36] caused by *mortalin* silencing in HeLa cells and the dopaminergic SH-SY5Y cell line. To address whether promoting autophagy is detrimental or protective *in vivo*, we at first modulated autophagic flux genetically. Quantification of larval locomotion revealed that pan-neuronal overexpression of *Atg1* at 25°C is sufficient to cause a sluggish righting reflex. Concomitant overexpression of elav>*mort*^*KK*^ did not further exacerbate this phenotype (Figure 6A). Using the Gal4/Gal80 system, we investigated the effect of *Atg1* overexpression on *Hsc70-5* knockdown related phenotypes in flies by performing the longevity assay (Figure 6B). Knockdown of *Hsc70-5* caused a significant reduction in median and maximum life expectancy compared to control flies. *Atg1* overexpression in the control background enhanced median but not maximum survival. Notably, concomitant overexpression of *Atg1* with *mort*^*KK*^ caused a reduction in both median and maximum lifespan compared to *mort*^*KK*^ alone (Figure 6B). *Atg1* overexpression in the elav>*mort*^*KK*^,tub-GAL80^ts^ background exacerbated the loss of *Hsc70-5* induced climbing impairment (Figure 6C); however, a slight improvement was observed in wing phenotypes (Figure 6D). *Atg1* overexpression alone did not impair adult locomotion of 4-day-old flies examined using the climbing ability nor induce abnormal wing phenotypes (Figure 6C and D). These results suggest that overexpression of *Atg1* did not modify *Hsc70-5* knockdown induced defects at the larval stage but reduced lifespan and climbing ability in flies.

**Figure 6.**
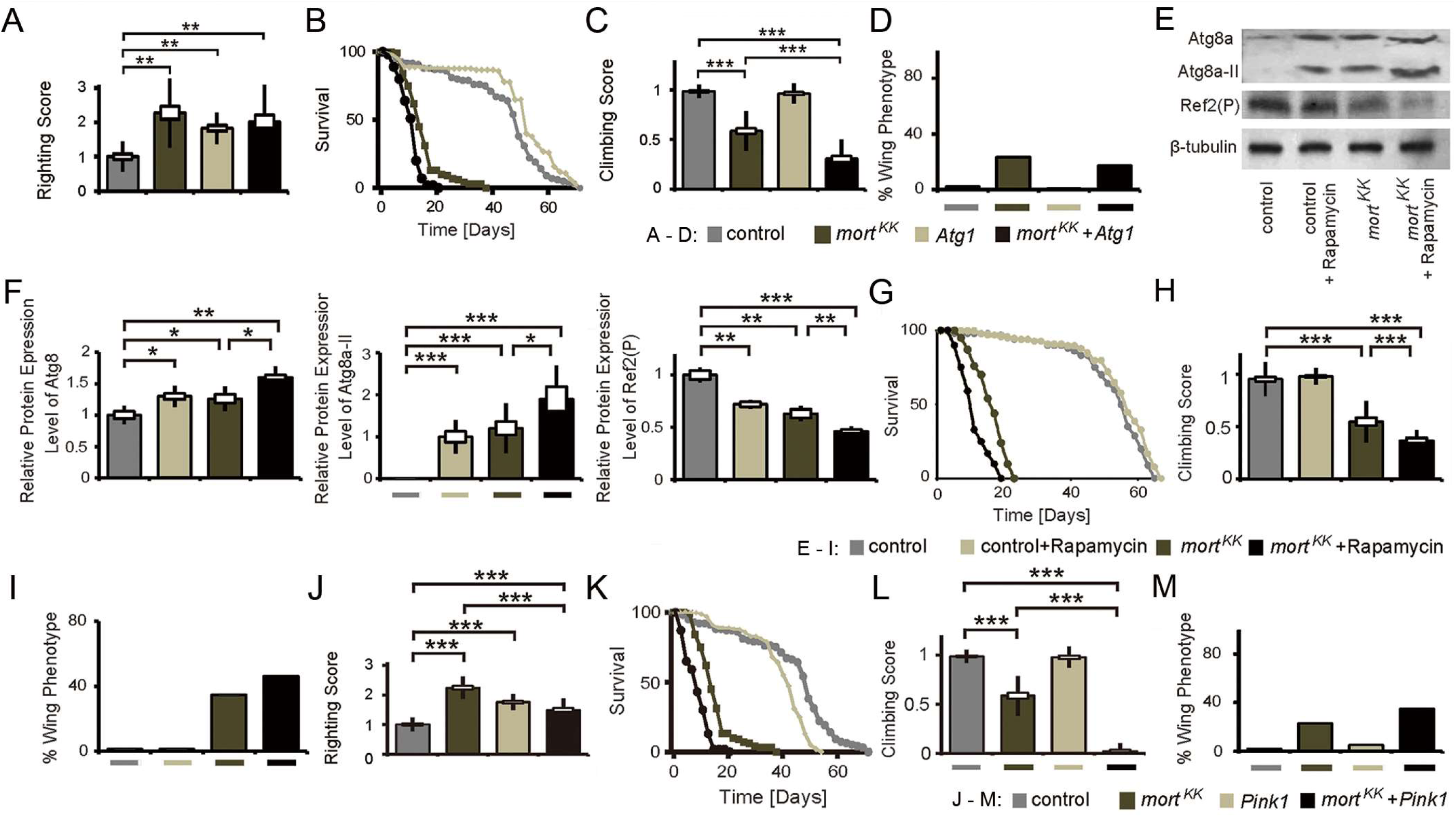
Autophagy induction did not rescue symptomatic phenotypes in *Hsc7-5/mortalin* knockdown flies. (**A**) Quantification of larval righting reflex, (**B**) Lifespan of flies at 25°C, (**C**) Climbing ability of 4-day-old flies and (**D**) Percentage of flies with defective wing phenotype expressing *Atg1* in the elav>*mort*^*KK*^,tub-GAL80^ts^ background. (**E**) Western blot showing levels of Atg8a, Atg8a-II, and Ref2(P) in rapamycin-treated flies in the elav>*mort*^*KK*^,tub-GAL80^ts^ background. β-tubulin was used as a control. (**F**) Quantifications of protein levels of Atg8a, Atg8a-II, and Ref2(P). (**G**) The lifespan of flies with rapamycin treatment in the elav>*mort*^*KK*^,tub-GAL80^ts^ background at 25°C. (**H**) The climbing ability of 4-day-old flies, and (**I**) Percentage of flies with defective wing phenotype with rapamycin treatment in the elav>*mort*^*KK*^,tub-GAL80^ts^ background. (**J**) Quantification of larval righting reflex, (**K**) Lifespan of flies at 25°C, (**L**) Climbing ability of 4-day-old flies, and (**M**) Percentage of flies with defective wing phenotype expressing *Pink1* in the elav>*mort*^*KK*^,tub-GAL80^ts^ background. * p<0.05, ** p<0.01, *** p<0.001.

Rapamycin has been used to induce autophagy in *Drosophila* [37]. Feeding rapamycin is sufficient to enhance autophagy as demonstrated by increased accumulation of autophagy markers Atg8a and relative levels of Atg8a-II, in addition to reduced levels of Ref(2)P, homolog of mammalian p62 (Figure 6E and F). *Hsc70-5* knockdown caused the same effects in fly heads (Figure 6E and F). We investigated whether supplementing rapamycin could further enhance autophagy caused by *Hsc70-5* knockdown. Quantification of western blots showed a further increase in the accumulation of total Atg8a and relative levels of Atg8a-II with concomitant reduction of Ref(2)P (Figure 6E and F). Rapamycin treatment prolonged lifespan in control flies but caused a reduction in elav>*mort*^*KK*^,tub-GAL80^ts^ flies (Figure 6G). Moreover, rapamycin treatment exacerbated the climbing impairment and wing phenotypes in symptomatic flies (Figure 6H and I).

*PINK1* has been implicated in familial PD. It regulates the degradation of old and dysfunctional mitochondria in health and upon exposure to the mitochondrial uncoupler CCCP [6-8,38,39]. Hence, we examined if *Pink1* overexpression could modulate the loss of *Hsc70-5* phenotypes. *Pink1* overexpression was sufficient to induce a sluggish righting reflect at the larval stage (Figure 6J). In elav>*mort*^*KK*^ larvae it rescued locomotion defects (Figure 6J). Ectopic *Pink1* expression in the elav>*mort*^*KK*^,tub-GAL80^ts^ genetic background reduced lifespan (Figure 6K) and exacerbated the climbing impairment and wing phenotypes in symptomatic adult flies (Figure 6L and M).

### *Atg1* knockdown rescued synaptic mitochondrial loss and synaptic defects observed in symptomatic larvae

*Atg1* is required for the initiation of autophagosome formation [40]. We thus characterized the cellular changes caused by modulation of *Atg1* expression in the elav>*mort*^*KK*^ background. *Atg1* knockdown reversed the loss of mitochondria observed at the NMJs of elav>*mort*^*KK*^ larvae (Figure 7A) and rescued the decrease in mitochondria area fraction, mitochondrial number, size, and altered morphology at the NMJ in elav>*mort*^*KK*^ larvae (Figure 7B).

**Figure 7.**
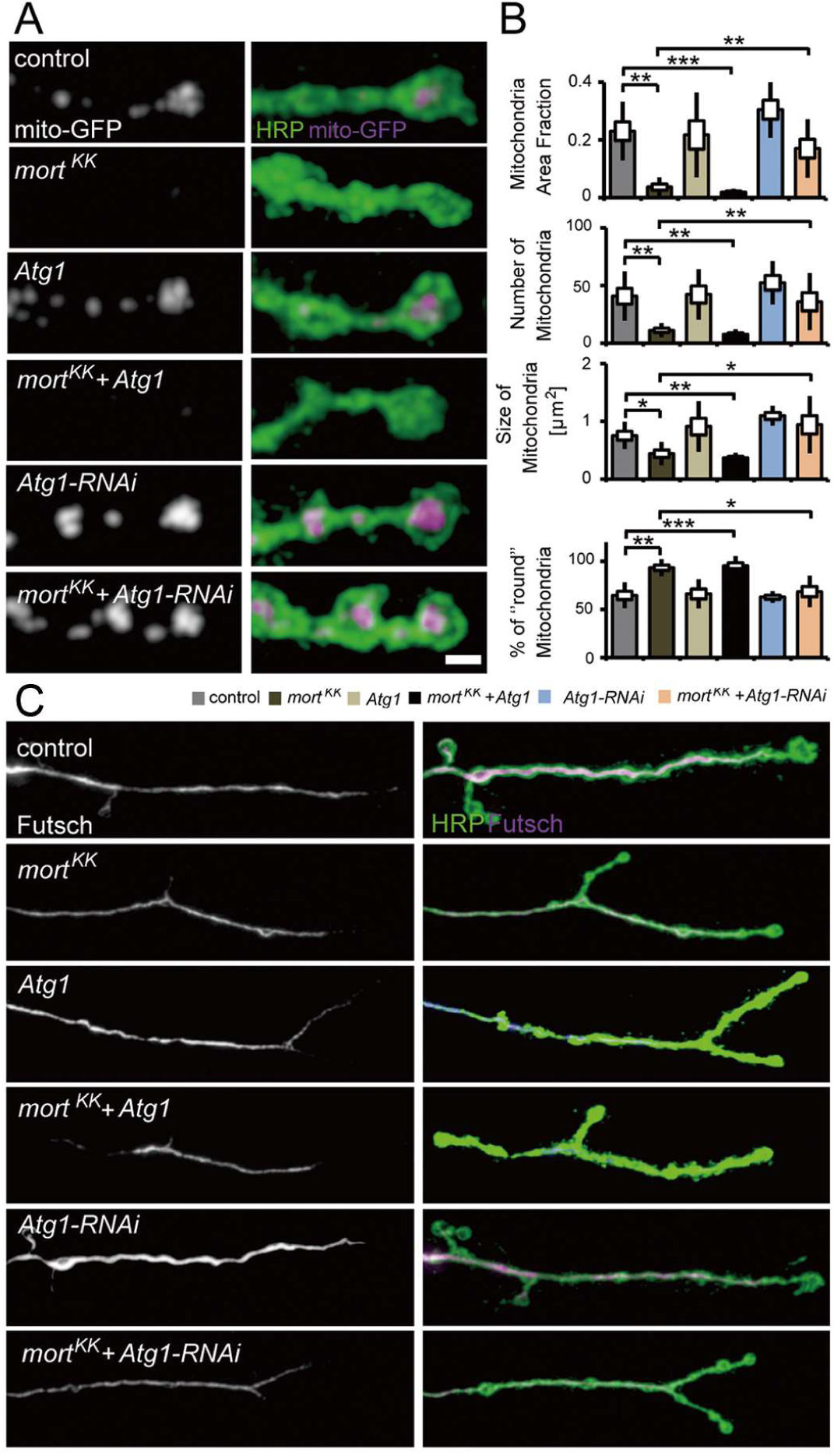
Autophagy inhibition alleviated mitochondrial and MT defects caused by *Hsc70-5/mortalin* knockdown. (**A**) Confocal images of larval NMJ expressing mito-GFP (magenta) labeled with HRP (green). Scale bar: 2 μm. (**B**) Quantification of mitochondrial area fraction, number, size, and morphology. (**C**) Confocal images of NMJ labeled with HRP (green) and Futsch (magenta). Scale bar: 10 μm. * p<0.05, ** p<0.01, *** p<0.001.

NMJs of early symptomatic elav>*mort*^*KK*^ larvae were characterized by reduced presence of SV marker proteins and Az components, impaired synapse maturation, and alterations in the MT cytoskeleton. We next assessed MT abundance as a marker for synapse stability. *Atg1* knockdown restored usual MT abundance in elav>*mort*^*KK*^ larvae (Figure 7C). Furthermore, *Atg1* knockdown in elav>*mort*^*KK*^ larvae alleviated impairments in VGlut (Figure 8A and B) and Brp abundance (Figure 8C and D), and synapse maturation impairment (Figure 8E and F). Overexpression of *Atg1* failed to rescue defects in MT abundance (Figure 7C), SV protein abundance (Figure 8A and B), and synaptic maturation defects caused by loss of *Hsc70-5* at the NMJ (Figure 8E and F). These findings indicate that restoring mitochondrial mass by *Atg1* knockdown could rescue loss of *Hsc70-5* induced synaptic defects.

**Figure 8.**
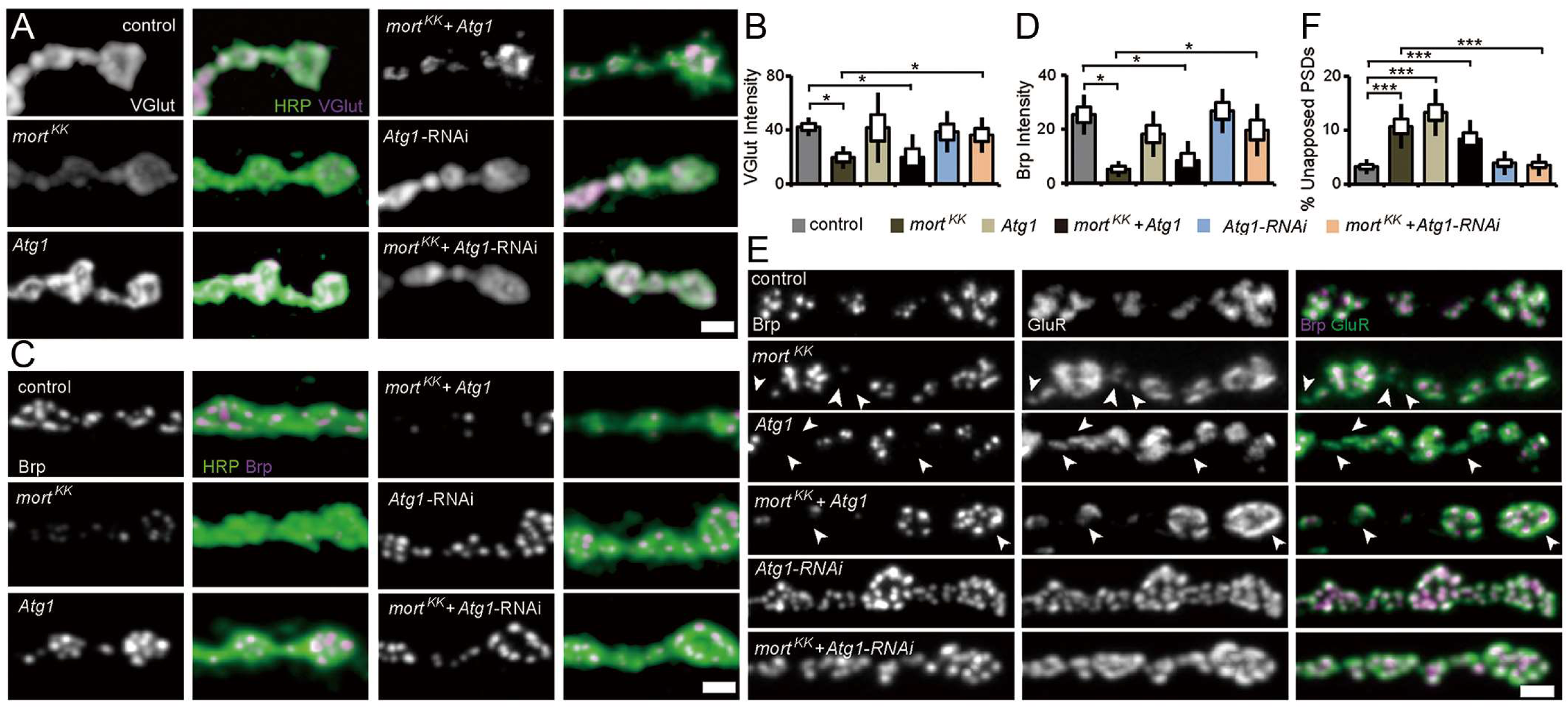
Autophagy inhibition alleviated synaptic defects caused by *Hsc70-5/mortalin* knockdown. (**A, C**) Confocal images of larval NMJ labeled with HRP (green), VGlut, and Brp (magenta). Scale bar: 2 μm. (**B, D**) Quantification of VGlut and Brp level at NMJ following *Atg1* ectopic expression and knockdown in elav>*mort*^KK^ background. (**E**) Confocal images of larval NMJ labeled with Brp (magenta) and GluR (green). Arrowheads pointing out regions where presynaptic Brp labels were not detected in PSDs. Scale bar: 2 μm (**F**) Quantification of unapposed glutamate receptor fields. * p<0.05, *** p<0.001.

### Autophagy suppression was protective against oxidative stress

Next, we tested whether knockdown of *Hsc70-5* enhances vulnerability to oxidative stress. We treated young flies with hydrogen peroxide (H_2_O_2_). H_2_O_2_ induces generalized oxidative stress which is less specific than the effects of the mitochondrial complex inhibitors like rotenone or paraquat [41]. *DJ-1* and *LRRK* are crucial for protection against oxidative stress [41,42]. H_2_O_2_ has also been shown to reduce the lifespan of flies that display a reduced function of the PD-associated proteins *DJ-1* and *LRRK*. elav>*mort*^*KK*^,tub-GAL80^ts^ flies were more vulnerable to H_2_O_2_ treatment than control flies (Figure 9A). While *Atg1* knockdown caused a minor reduction of lifespan upon exposure to oxidative stress compared to control, it proved to be effective in restoring the diminished stress resistance in the elav>*mort*^*KK*^,tub-GAL80^ts^ background (Figure 9A). We thus conclude that suppression of autophagy in the elav>*mort*^*KK*^,tub-GAL80^ts^ background might be beneficial for survival under oxidative stress.

**Figure 9.**
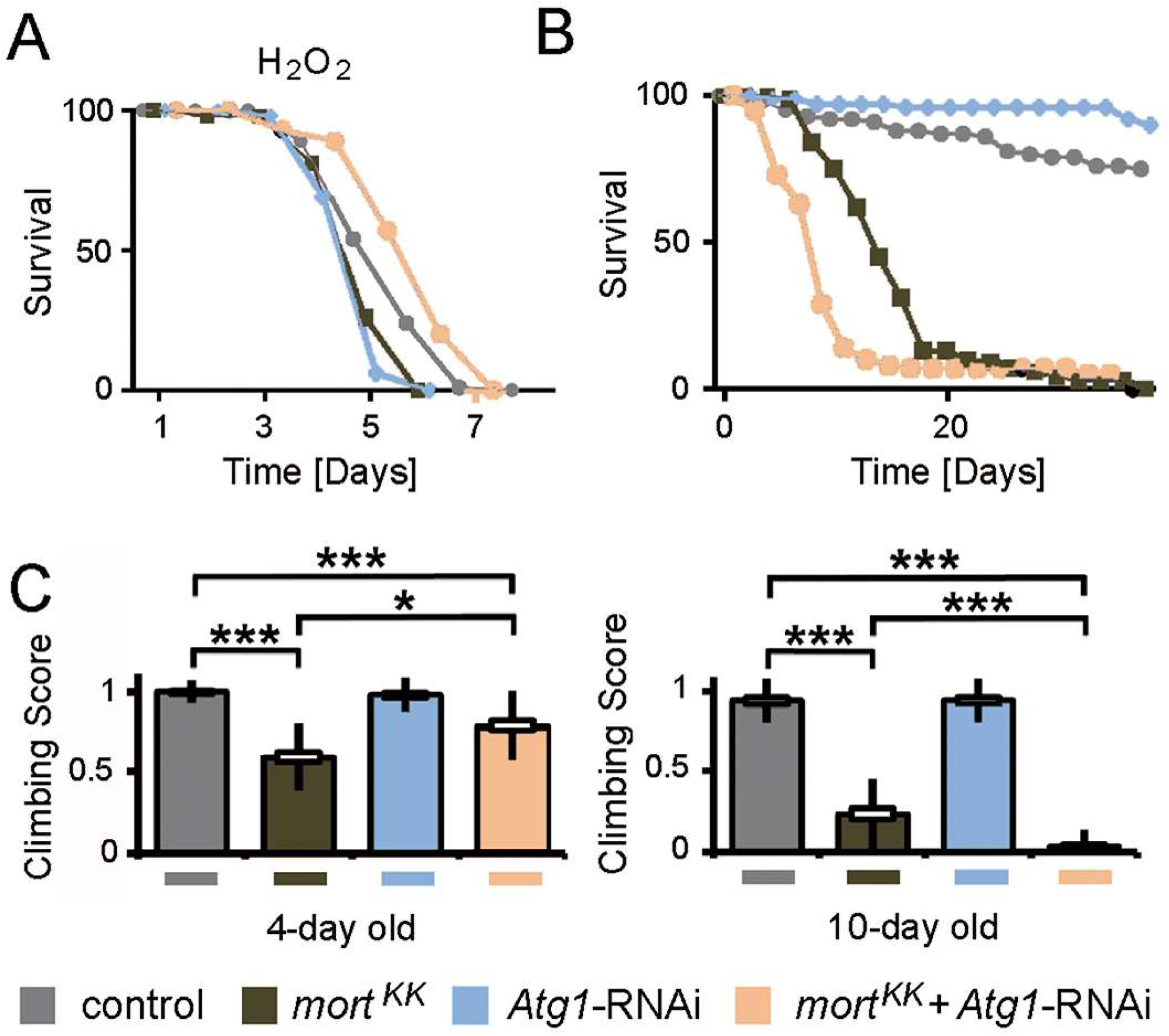
Concomitant *Atg*1 and *Hsc70-5/mortalin* knockdown rescued longevity under oxidative stress but was detrimental upon aging in baseline conditions. (**A**) Lifespan of flies at 25°C following induction of generalized oxidative stress. Flies were fed with 5% hydrogen peroxide sucrose solution. (**B**) Lifespan of flies at 25°C on the standard diet. (**C**) The climbing ability of 4- and 10-day-old flies expressing *Atg1* knockdown in the elav>*mort*^*KK*^,tub-GAL80^ts^ background. * p<0.05, *** p<0.001.

### Autophagy suppression improved healthspan but did not extend lifespan

To investigate the long-term impact of *Atg1* knockdown, we examined the lifespan of flies under normal conditions (Figure 9B). *Atg1* knockdown led to a reduction of the lifespan in elav>*mort*^*KK*^,tub-GAL80^ts^ flies. As a minor decrease in lifespan had been observed, we thought to assessed the effect of *Atg1* knockdown at various pathological stages. *Atg1* knockdown was beneficial in 4-day old symptomatic flies and improved locomotion. In 10-day old flies, concomitant knockdown resulted impairment in climbing using a less challenging (Figure 9C). We referred to these 10-day old flies as late-symptomatic flies (Table S1). We conclude that reduced autophagy flux is beneficial for improving locomotion ability in *Hsc70-5* reduced background in young flies, but impairs climbing ability and lifespan in elav>*mort*^*KK*^,tub-GAL80^ts^ flies in the long run.

## Discussion

### Investigating the impact of rare *Hsc70-5/mortalin* mutations *in vivo*

Three mutations have been identified in *mortalin* in a congenital disease termed epiphyseal, vertebral, ear, nose, plus associated finding) syndrome or EVEN-PLUS [28]. Biallelic mutations were identified in three individuals from two families, including the previously reported R126W variant [17]. Functional studies investigating the impact of variant *HSPA9*^P509S^ revealed a lower ATP catalysis rate and reduced chaperoning activity compared to *HSPA9*^WT^ [17]. Next, morphological studies in human fibroblasts from a heterozygous carrier of variant *HSPA9*^A476T^ showed mitochondrial defects compared to *HSPA9*^WT^ homozygous sibling [17]. Another study investigated the effects of human variant R126W, and P509S in yeast by generating analogous substitutions in the Saccharomyces cerevisiae ortholog of *Hsc70-5*, the *SSC1* gene [16]. The substrate-binding domain mutation *SSC1* A453T was shown to cause mitochondrial dysfunction, enhanced ROS levels, and reduced ability to prevent aggregate formation of unfolded substrates [16].

In this study, we used a *Drosophila* model of loss of *Hsc70-5/mortalin* function to test the functional relevance of rare *mortalin* mutations *HSPA9*^R126W^, *HSPA9*^A476T,^ and *HSPA9*^P509S^ *in vivo*. Overexpression of human and *Drosophila mortalin (HSPA9 and Hsc70-5)*, in a genetic background with reduced *Hsc70-5*, unlike mutant variants *HSPA9*^R126W^, *HSPA9*^A476T^ and *HSPA9*^P509S^, were able to rescue the loss of synaptic mitochondria, locomotion defects, abnormal wing phenotype, and reduced ATP levels. This study provides evidence that the investigated amino acid replacements in the ATPase and substrate-binding domain impair Mortalin function. *In vitro* overexpression studies have suggested no difference in overexpression levels or localization of the mutant variants [17] and future studies should address this in *in vivo* setting.

### *Hsc70-5/mortalin* knockdown induced synaptic defects

Neuronal mitochondria have been implicated in diverse functions including spine formation [43], synaptic plasticity [44], and axonal branching [45]. Loss of *Miro* dramatically excludes mitochondria from distal compartments and causes a gradual time-dependent reduction in EJP amplitude following 10Hz stimulation, although no changes were observed in EJP or mEJP amplitude under baseline conditions [24]. Similar findings were reported in *Drp1* mutants where dramatic synaptic mitochondria reduction was observed [46].

Loss of neuronal mitochondria as a consequence of *Hsc70-5* silencing causes a reduction in EJP and mEJP amplitudes and quantal content (Figure 3A-C). There was an increase in the frequency of mEJP events (Figure 3C), possibly due to the elevation of intracellular calcium levels following loss of *Hsc70-5*. More frequent synaptic failure was observed upon stimulation at 10Hz (Figure 3F). Failed responses that are caused by the depletion of synaptic vesicles are preceded by a gradual decline in EJP amplitude [46]. However, in our case, the amplitude of the remaining EJPs was not affected. Thus, failures might instead be caused by impaired propagation of action potentials as described for *Atpα* mutant [32].

Cellular analysis of larval NMJ in *Hsc70-5* knockdown animals did not reveal gross impairments in the morphology of distal compartments (Figure S2) or inhomogeneity of presynaptic membrane or synaptic footprints (Figure 3B) that are biomarkers for early and late stages of synapse disassembly, respectively [25,26]. Importantly, defects in the availability of synaptic components, such as CSP, VGlut, and Brp (Figure 1), were observed at a stage in which no major [47] neurodegeneration was noted. Hence, the protective effects of blocking mitophagy described in this study were likely related to mitochondrial function rather than susceptibility to cell death. At this stage of disease progression, RNAi against *Atg1* was sufficient to reverse defects in development, regulation of synaptic vesicles, synaptic terminal stability, and function (Figures 7 and 8). Furthermore, *Atg1*-RNAi restored mitochondria abundance. In addition, it also reversed the only putatively degenerative change we found in Hsc70-5 knockdown larvae: MT cytoskeletal alterations (Figure 7B).

### Epistatic interaction of *Hsc70-5/mortalin* with the autophagic machinery

Mitophagy, a type of macro-autophagy that targets specifically mitochondria, is increased upon partial loss of *mortalin* function in *Drosophila* and human fibroblasts [14]. Increased autophagic flux can be likened to a double-edged sword that protects neurons from chronic oxidative stress but might accelerate mitochondrial loss under conditions of premature mitophagy or impaired mitochondrial biogenesis [13,48]. The beneficial effects of *Parkin* overexpression in mouse embryonic fibroblasts treated with siRNA against *mortalin* were dependent on intact autophagic machinery, suggesting a potential therapeutic benefit of upregulating autophagy [17]. However, the interpretation of suppressed apoptosis in HEK293 cells and tumor-derived SH-SY5Y cells is complicated by the fact that (like most tumor-derived culture cells) these cells primarily generate ATP via aerobic glycolysis and are therefore not dependent on oxidative phosphorylation [49]. Thus, the balance between costs and benefits of effective mitochondria removal might differ *in vivo*.

We previously demonstrated that *Hsc70-5* knockdown in *Drosophila* caused a severe loss of synaptic mitochondria and cellular ATP depletion [13]. The *Drosophila* NMJ has proved to be a particularly useful model as it is easily accessible and morphologically complex. These characteristics allowed us to model a highly active synaptic population that is vulnerable to impairment in intracellular trafficking because it is located very distantly to the neuronal soma. The *in vivo* data obtained in this study did not provide evidence for the therapeutic benefit of promoting autophagic flux following impaired mitochondrial function. Ectopic *Atg1* expression, which was sufficient to induce autophagy in *Drosophila* [50] did not rescue impaired locomotion or mitochondrial mass following *Hsc70-5* knockdown in the symptomatic larval model (Figure 7A). Besides, *Atg1* overexpression did not reverse alterations in synaptic development at the larval NMJ (Figure 8). In symptomatic flies, *Atg1* overexpression exacerbated climbing defects and shortened lifespan caused by loss of *Hsc70-5*. The inability of *Atg1* to exacerbate *Hsc70-5* knockdown associated mitochondrial abundance or cellular defects at the larval stage might be explained by the fact that overexpression of *Atg1* alone does not reduce mitochondrial abundance (Figure 7A). It is also possible that the loss of mitochondria upon *Hsc70-5* knockdown is so extreme that further *Atg1* overexpression is unable to exacerbate the already severe defect (Figures 7 and 8).

The analysis of *Atg1* overexpression in larval stages was complicated by the fact that *Atg1* overexpression alone caused a sluggish righting phenotype (Figure 6A). *Pink1* overexpression alone also caused a sluggish righting phenotype in larvae and reduced life expectancy in adult flies (Figure 6). Nevertheless, both *Atg1* and *Pink1* overexpression exacerbated loss of *Hsc70-5* associated defects in locomotion and longevity in adult flies. Future studies might address the interaction between *Hsc70-5* and regulators of mitochondrial quality control such as *Pink1* and *parkin* and proteins involved in mitochondrial dynamics in more detail.

Using a functional genetic screen we identified that knockdown of autophagy-related genes in elav>*mort*^*KK*^,tub-GAL80^ts^ flies rescued climbing defects, abnormal wing posture, and ATP levels in 4-day old flies (Figure 5B). However, knockdown of autophagy was detrimental in late-symptomatic 10-day old flies.

How can these data be reconciled with reports suggesting that increased autophagy might be of therapeutic potential in PD [51] ? Mitochondrial quality control mechanisms are very divergent. In particular, the molecular mechanisms underlying mitochondrial stress responses in *Drosophila* and mammals might differ. While neither *Parkin* nor *Pink1* knockout-mice display noticeable behavioral or morphological changes under baseline conditions, loss of *parkin* or *Pink1* in *Drosophila* is detrimental and results in decreased lifespan and apoptotic flight-muscle degeneration [6,7,52,53]. The mammalian system might be equipped with more elaborate compensatory mechanisms and might display more functional redundancy, whereas flies might be particularly vulnerable to overactivation of mitophagy as evident by the extreme depletion of mitochondria from synapses (Figure 1B). Similar pathological induction of autophagy has been reported for hypoxic-ischemic brain injury and two toxin-based PD models in which blockade of autophagy proved to be protective [54].

### Stimulation of autophagy under conditions of oxidative stress

Oxidative and nitrosative stress is associated with various neurodegenerative diseases, including PD [55,56]. Pathogenic mechanisms include excitotoxicity, endoplasmic reticulum stress, protein aggregation, and damage to mitochondria, which are generators and targets of ROS. Flies affected by *Hsc70-5* loss displayed increased vulnerability to oxidative stress (Figure 9A). These effects might be mediated through the Mortalin interaction partner DJ-1 [57]. DJ-1 associated with autosomal-recessive early-onset familial PD is important for response against oxidative stress [58,59]. Loss of DJ-1 renders cells vulnerable to oxidative stress [57,60,61]. Notably, the interaction between Mortalin and DJ-1 is vital for the control of oxidative stress in hematopoietic stem cells [57].

Cross-talk between autophagy and ROS/nitrosative stress in the context of cell signaling and pathological protein damage is considered a significant obstacle for the therapeutic modulation of autophagy [62,63]. Thus, it is noteworthy that suppression of autophagy was beneficial in the context of combined mitochondrial dysfunction and oxidative stress in the symptomatic adult model (Figures 7 and 9) presented in this study. The *in vivo* evidence presented here suggests that reduced rates of autophagy might be protective for neurons compromised by pathologically increased levels of mitophagy. However, we observed that detrimental side effects exceeded protective benefits in the long-term (Figure 9).

## Materials and Methods

### Molecular biology

cDNA constructs for *HSPA9*^*WT*^, *HSPA9*^*R126W*^, *HSPA9*^A476T^, *and HSPA9*^*P509S*^ were received from Rejko Krüger (University of Luxembourg). The constructs were recloned and inserted into a modified pUAST attB vector (using BamH I and Xho I). The full-length *Drosophila Hsc70-5* (GM13788) was inserted into a modified pUAST attB vector. Details of fly strains have been provided in the supplementary information (Materials and Methods, Table S2).

### Bar charts

The standard error of mean (SEM) and standard deviation (SD) are shown as a box and black line.

### Electrophysiology

Electrophysiological recordings were performed essentially as previously described [19,25].

### Staining and imaging larval neuromuscular junctions

Dissection and labeling of size-matched mid-3^rd^ instar larvae were performed using previously described protocols [25]. For more information on antibody source and concentrations, refer to supplementary information.

### Fat body assay for investigating autophagy

Assay to validate functionality of UAS-*Atg1* overexpression or RNAi constructs against autophagy related proteins were performed using a mosaic genetic analysis in larval fat body cells essentially as previously described [34,35]. Refer to supplementary information for details.

### ATP measurements

ATP levels in head homogenates were measured and normalized using a luciferase-based bioluminescence assay as described earlier [13]. Five heads of female flies were homogenized in 6M guanidine-HCl and frozen in liquid nitrogen. Next, samples were boiled for 3 min, cleared by centrifugation at 14,000g for 5 min, and diluted to measure protein concentration (1:10 diluted samples, Bradford Assay Kit, Sigma-Aldrich, B6916) and ATP level (1:2,000 diluted samples, ATP Determination Kit Sensitive Assay, Biaffin GmbH & Co KG, LBR-P010). ATP levels were normalized to the protein concentration.

### Locomotion analysis

Climbing assays were, unless otherwise noted, conducted as previously described [14]. Climbing male flies was monitored by analyzing their performance to climb 6 cm (challenging assay, used in Figure 4D or 5D) or 3 cm (less challenging assay used in Figure 9C for 10-day old flies) within 14s. A successful attempt was scored as 1, and failure to reach the top as 0. Each fly was assessed three times to calculate the average climbing score. At least 40 flies per genotype were analyzed. Larval locomotion was investigated by examining larvae crawling speed and righting assay. Larvae locomotion speed was quantified as previously described [25].

### Longevity assay

*Drosophila* larvae were transferred from 18°C to 25°C to boost UAS-transgene expression 5 days AEL. Adult male flies were maintained at 25°C in groups of 20 or fewer. 100 flies were assayed per genotype. No anesthesia was used in survival experiments.

### Rapamycin treatment

Rapamycin (LC Laboratories, 53123-88-9) was dissolved in ethanol and added to standard fly food at appropriate concentrations (0.1 and 200 μM). For control food (0 μM), ethanol alone was added. 5 days AEL, larvae were transferred from 18°C to 25°C to induce transgene expression on 0 μM (Control) and 0.1 μM (Rapamycin) food. Newly hatched male flies were continually maintained in 0 μM (Control) and 200 μM (Rapamycin) food at 25°C for longevity assay.

### Western blot

Western blot analysis was performed on 20 µg samples of 4-day-old fly head protein using antibodies for Atg8a, Ref2(P) (rabbit polyclonal, 1/1000, a gift from Gábor Juhász, Eotvos Lorand University) and β-tubulin (mouse monoclonal, 1/1000, Developmental Studies Hybridoma Bank) using chemiluminescent detection.

## Acknowledgement

We thank the Developmental Studies Hybridoma Bank maintained by the University of Iowa for antibodies. We are grateful to the Bloomington and VDRC Stock Centers as well as all researchers not mentioned explicitly who made their stocks available via the Bloomington Stock Center. We thank Raphael Zinser for excellent assistance. We thank the DKFZ Light Microscopy Facility. We thank the Fritz Thyssen Foundation (grant 10.12.1.192 awarded to TMR), the Hertie Foundation (for stipends awarded to JZ, SBH, NV), the Chica and Heinz Schaller Foundation (for supporting SBH, NMD, TRJ), and Max Planck Society (for supporting TMR).

## List of abbreviations

AEL: after egg laying
AZ: active zone
Brp: Bruchpilot
CSP: cysteine string protein
DLG: discs large
eEJPs: evoked excitatory junctional potentials
GluR: glutamate receptor
mEJP: miniature excitatory junctional potentials
MT: microtubule
NMJ: neuromuscular junction
PD: Parkinson disease
*Pink1*: *PTEN-induced putative kinase 1*
PSD: postsynaptic density
SSR: subsynaptic reticulum
SV: synaptic vesicle
Vglut: vesicular glutamate transporter

## Supplementary information

### Materials and methods

#### Fly strains

Transgenic fly stocks were obtained either from Indiana University Stock Center (Bloomington, IN, USA) or Vienna *Drosophila* RNAi center unless otherwise noted. Transgenic stocks UAS-*HSPA9*^*WT*^, UAS-*HSPA9*^*R126W*^, UAS-*HSPA9*^*A476T*^, UAS-*HSPA9*^*P509S*,^ and UAS-*Hsc70-5* were created by BestGene using integrase mediated site-specific transgenesis at cytological position 68A4 (Fly strain BDSC 8622).

#### Fly culture conditions

Flies were raised on standard cornmeal/agar medium. To circumvent pupal lethality caused by *Hsc70-5/mortalin* knockdown and to analyze behavioral defects in adult flies, we utilized the Gal4/Gal80 system (elav>*mort*^*KK*^,tub-GAL80^ts^). This allowed us to achieve late-onset conditional knockdown, by raising larvae at 18°C before transferring them to 25°C at 5 days after egg laying (AEL). Flies were kept at 25°C during development for analysis of wing phenotype, climbing defects, longevity, ATP levels from heads, and temperature-induced paralysis at appropriate ages.

#### Morphological analysis

Analyses of mitochondria and NMJs were performed as previously described [1]. The area fraction occupied by mitochondria was used as the index of mitochondrial mass. To quantify Futsch loops, the number of loops located to the two most distal boutons at each terminal of the NMJ was scored. The total number of loops in these regions was normalized to the number of terminals.

#### Staining and imaging larval neuromuscular junctions

Larvae expressing GFP were fixed for 3 min (4% paraformaldehyde in phosphate-buffered saline (PBS). Goat anti-horseradish peroxidase (HRP)-Cy3 antibody was obtained from Dianova (Hamburg, Germany). Primary antibodies were used in the following dilutions: mouse anti-Bruchpilot (Nc82) (1:100), mouse anti-CSP (DSCP-2) (1:150), rabbit anti-DVGlut (1:1000), mouse anti-Dlg (1:100), and mouse anti-Futsch (22C10) (1:100). Confocal imaging of NMJ 4 at Segment A5 of mid-3^rd^ instar larvae was performed using a Zeiss LSM 710 microscope (Carl Zeiss, Oberkochen, Germany) using a 40× Plan-Apochromat 1.4 N.A. oil objective essentially as previously described (unless otherwise noted described) [2-4]. A voxel dimension of (x/y/z) 100×100×500 nm was utilized. The pinhole size was 1 Airy Disc. Images were scaled by a factor of 2 before Gaussian blur filtering was applied (pixel radius=2). Gamma values were set to 0.75. For quantitative comparisons of intensities, standard imaging settings were chosen that avoided oversaturation. ImageJ Software Version 1.43e (National Institutes of Health, Bethesda, MD, USA) was used for image processing.

#### Larval fat body assay for investigating autophagy

Starvation of 2^nd^ instar stage larvae was performed for 5-6 hours in fresh empty vials on a filter paper soaked with H_2_O. For imaging, 2^nd^ instar larvae that were well fed or starved for 4-5 hours were cut open and turned inside out like a ‘sock’ before removal of fat bodies. Fat body tissues were stained with Dapi for 2 minutes, briefly washed in PBS, and mounted on a glass slide using Vectashield and imaged immediately using a Leica SP5 II confocal imaging system using a 40× Plan-Apochromat 1.4 N.A. oil objective. The entire procedure from fat body isolation to imaging was completed within 30 minutes.

#### Morphological analysis

Analyses of mitochondria and NMJs were, unless otherwise noted, performed as previously described [1]. The area fraction occupied by mitochondria was used as the index of mitochondrial mass. To quantify Futsch loops, the number of loops located to the two most distal boutons at each terminal of the NMJ was scored. The total number of loops in these regions was normalized to the number of terminals.

## Figure legends for supplementary figures

**Figure S1.**
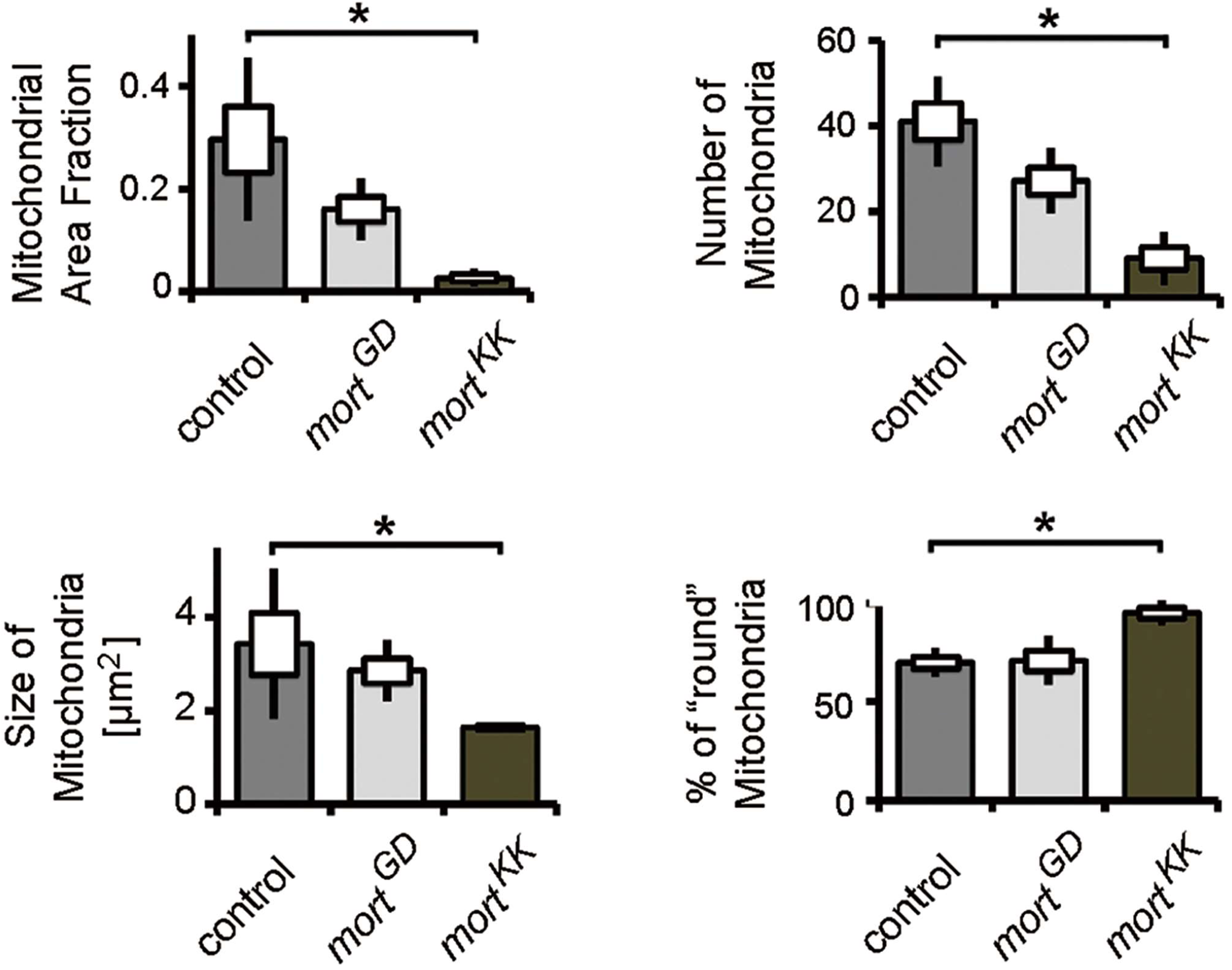
Quantification of mitochondrial parameters in *Hsc70-5/mortalin* knockdown larvae. Quantification of mitochondrial area fraction, number, size and shape in control, *mort*^GD^ and *mort*^KK^ larvae. Standard error of mean (SEM) and standard deviation (SD) were shown as a box and a black line, respectively. * p<0.05.

**Figure S2.**
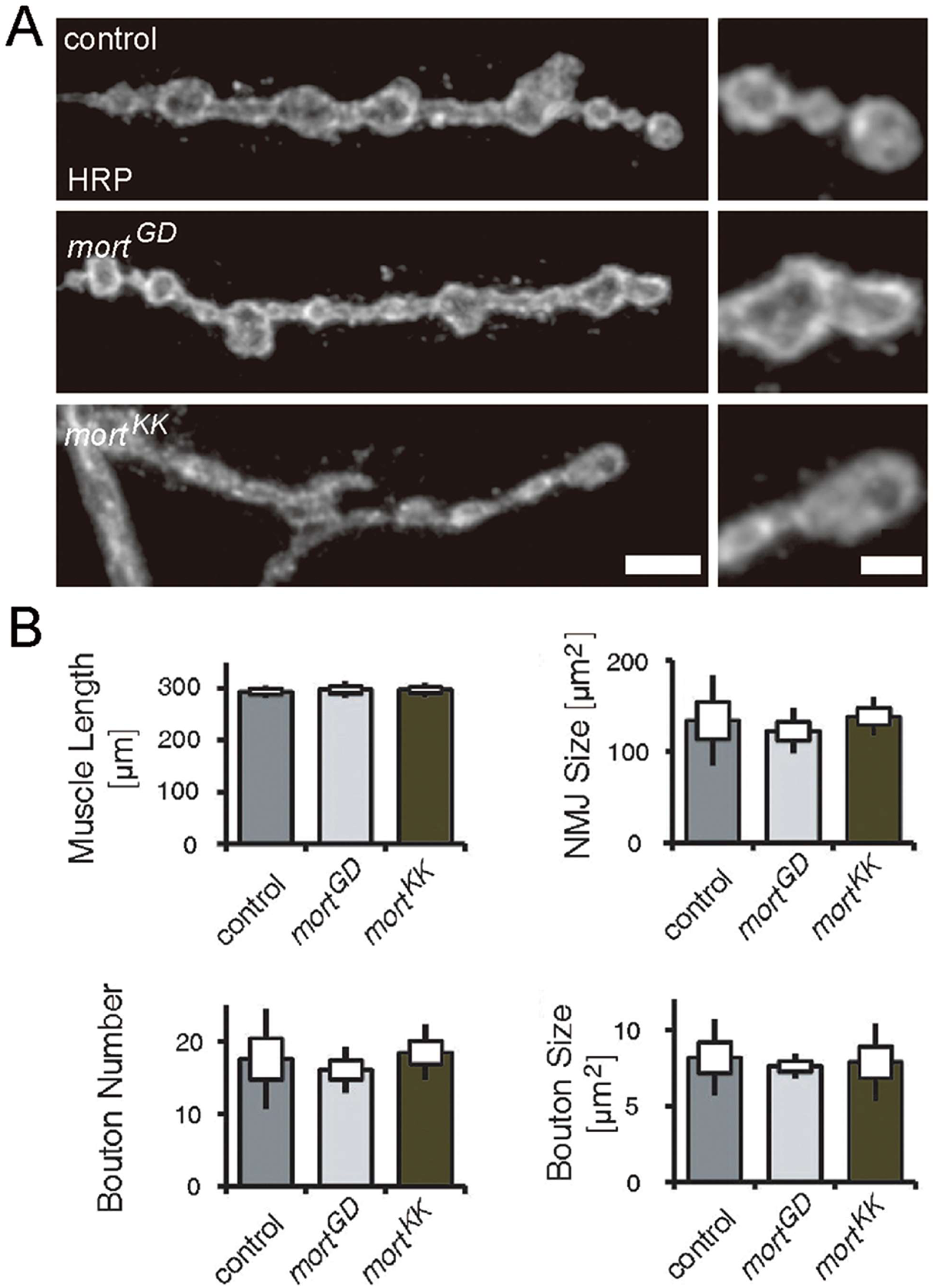
Quantification of synaptic terminals in *Hsc70-5/mortalin* knockdown larvae. (**A**) Confocal images of NMJ labeled with HRP-Cy3. Scale bar: 5 μm, Enlargement: 2 μm. (**B**) Quantification of muscle length, NMJ size, number, and bouton size of synaptic boutons in control, *mort*^GD^ and *mort*^KK^ larvae.

**Figure S3.**
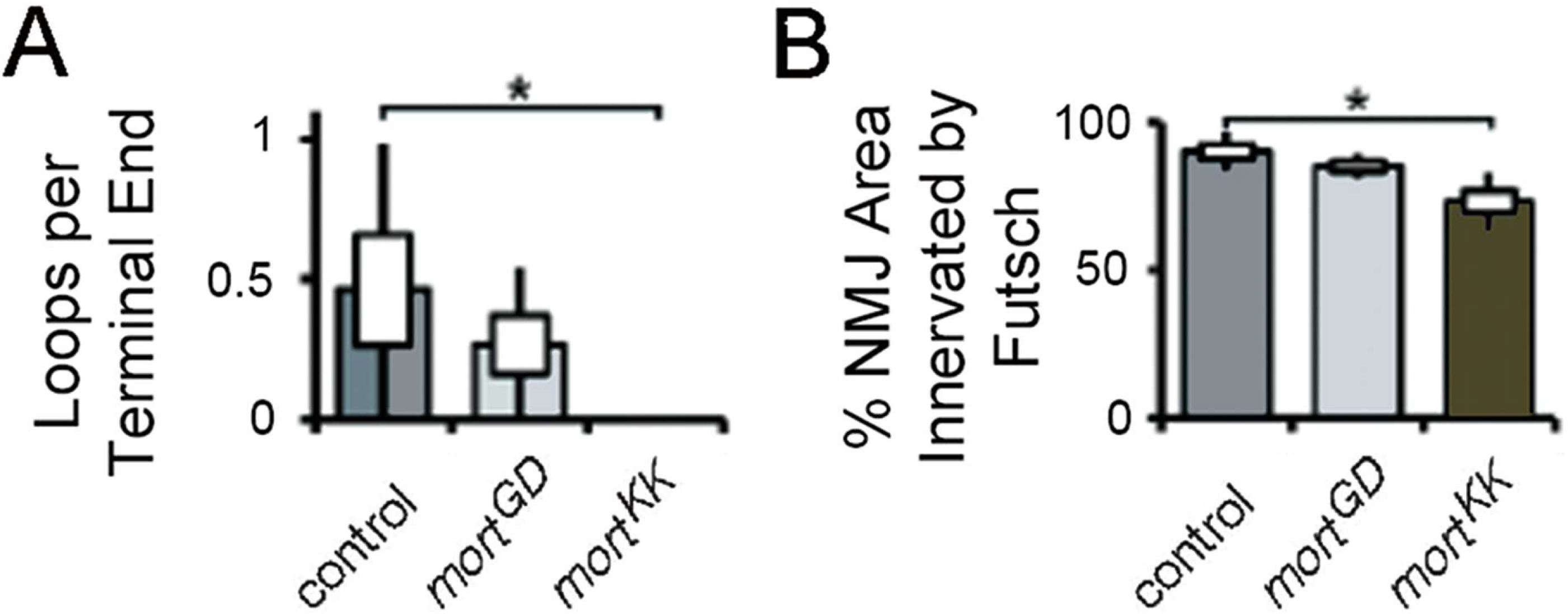
Quantification of Futsch staining in nerve terminals. (**A**) Futsch loops per terminal end and (**B**) % NMJ area innervated by Futsch in indicated genotypes. The standard error of mean (SEM) and standard deviation (SD) were shown as a box and a black line, respectively. * p<0.05.

**Table S1.** Categorization of pathological stages in larval and adult stages. (**1A**) Presymptomatic larvae: L3 elav>*mort*^GD^ larvae had no detectable locomotion defects compared to control and were categorized as presymptomatic. (**1B**) Symptomatic larvae: L3 elav>*mort*^KK^ larvae displayed lower crawling velocity and slower righting reflex compared to control and were classified as symptomatic. Similarly, elav>*mort*^KK^:tub-Gal80^ts^ L3 larvae were also classified as symptomatic due to a slower righting reflex compared to age-matched control larvae. (**2A**) Symptomatic adult: elav>*mort*^KK^:tub-Gal80^ts^ flies at 4 days were referred to as symptomatic based on their impaired climbing ability compared to control flies. (**2B**) Late-symptomatic larvae: At 10 days, elav>*mort*^KK^:tub-Gal80^ts^ flies displayed a more severe climbing defect compared to control and perform worse than 4-day old elav>*mort*^KK^:tub-Gal80^ts^. They were categorized as late-symptomatic.

| Stages | Model | Genotype/time point of analysis | Conditions | Locomotion phenotype |
| --- | --- | --- | --- | --- |
| Larva | (A) pre-symptomatic | elav>*mort*<sup>GD</sup>/L3 | 25°C | Crawling velocity and righting reflex unchanged |
|  | (B) symptomatic | elav>*mort*<sup>KK</sup>/L3 | 25°C | Crawling velocity lower and righting reflex slower |
|  |  | elav>*mort*<sup>KK</sup>,tub-Gal80<sup>ts</sup>/L3 | 1 day at 18°C → 25°C | Righting reflex slower |
| Adult | (A) symptomatic | elav>*mort*<sup>KK</sup>,tub-Gal80<sup>ts</sup>/5-day old | 5 days at 18°C → 25°C | Climbing efficiency reduced |
|  | (B) late-symptomatic | elav>*mort*<sup>KK</sup>,tub-Gal80<sup>ts</sup>/10-day old | 5 days at 18°C → 25°C | Exacerbated climbing defects |

**Table S2.** Fly strains. Fly strains utilized in this study and their sources.

| **Genes** | **CG number** | **Strain** |
| --- | --- | --- |
| *Atg1* | CG10967 | BL-26731 |
| *Atg101* | CG7053 | BL-34360 |
| *vsp15* | CG9746 | BL-35209 |
| *buffy* | CG8238 | BL-32060 |
| *Atg5* | CG1643 | BL-27551 |
| *Atg7* | CG5489 | BL-27707 |
| *Atg8* | CG32672 | BL-34340 |
| *Atg12* | CG10861 | BL-27552 |
| *SNAP25* | CG40452 | BL-27306 |
| *Sytβ* | CG42333 | BL-27293 |
| *Syx7* | CG5081 | BL-29546 |
| *Brp* | CG42344 | UAS-Brp-RNAi from Stephan Sigrist (Wagh et al., 2006) |
| *Vglut* | CG9887 | BL-27538 |
| *Rph* | CG11556 | BL-25950 |
| *Rop* | CG15811 | BL-28929 |
| *Sec15* | CG7034 | BL-27499 |
| *Endo A* | CG14296 | BL-27679 |
| *Eps-15* | CG16932 | BL-29578 |
| *Dj-1b* | CG6646 | BL-31261 |
| *LRRK* | CG5483 | BL-32457 |
| *p50* | CG8269 | BL-28596 |
| *p150* | CG9206 | BL-27721 |
| *tko* | CG7925 | BL-38251 |
| **Others** |  | **Strain** |
| *White RNAi* |  | VDRC-GD30033 |
| *mort RNAi* |  | VDRC-GD47745 |
| *mort RNAi* |  | VDRC-KK106236 |
| *Gal80<sup>ts</sup>* |  | BL-7019 |
| *UAS-Dicer* |  | BL-24646 |
| *UAS-Atg1* |  | Thomas Neufeld (University of Minnesota) |
| *hs-Flp;UAS-Dcr2;R4-mCherry-Atg8a,Act>CD2>Gal4,UAS-GFPnls* |  | mosaic stock from Gábor Juhász (Eotvos Lorand University) |
| *Gmr-Gal4* |  | Aaron Voigt (University Clinic Aachen) |
| *Lame duck-RNAi* |  | Aaron Voigt (University Clinic Aachen) |
| *UAS-Pink1* |  | Ming Guo (University of California Los Angeles) |

